# Effects of mid-frontal brain stimulation on sustained attention

**DOI:** 10.1101/641498

**Authors:** Martine R. van Schouwenburg, Ilja G. Sligte, Michael R. Giffin, Franziska Günther, Dirk Koster, Floortje S. Spronkers, Anna Vos, Heleen A. Slagter

**Author notes:** Corresponding author: Prof. Dr. Heleen Slagter, Department of Applied and Experimental Psychology, Vrije Universiteit Amsterdam, Van der Boechorststraat 7, 1081 BT, Amsterdam, the Netherlands.

## Abstract

Sustained attention is defined as the ability to maintain attention over longer periods of time, which typically declines with time on task (i.e., the vigilance decrement). Previous studies have suggested an important role for the dorsomedial prefrontal cortex (mPFC) in sustained attention. In two experiments, we aimed to enhance sustained attention by applying transcranial electrical current stimulation over the mPFC during a sustained attention task. In the first experiment, we applied transcranial direct current stimulation (tDCS) in a between-subject design (n=97): participants received either anodal, cathodal, or sham stimulation. Contrary to our prediction, we found no effect of stimulation on the vigilance decrement. In the second experiment, participants received theta and alpha transcranial alternating current stimulation (tACS) in two separate sessions (n=47, within-subject design). Here, we found a frequency-dependent effect on the vigilance decrement, such that contrary to our expectation, participants’ performance over time became worse after theta compared to alpha stimulation. However, this result needs to be interpreted with caution given that this effect could be driven by differential side effects between the two stimulation frequencies. To conclude, across two studies, we were not able to reduce the vigilant decrement using tDCS or theta tACS.

## Introduction

Sustained attention, the ability to filter incoming sensory information and maintain attention to this information for longer time periods, is an important aspect of cognitive functioning. Most people will experience that it is much easier to maintain attention over a longer period of time when in an engaging environment (for example driving a car in a city), compared to when they find themselves in a rather uneventful environment (driving a car on an empty highway). Indeed, research has shown large decrements in performance over time on sustained attention tasks that require participants to monitor and detect infrequent targets in a stream of non-targets [1,2]. Yet, sustained attention is crucial in daily life and in particular in jobs such as air traffic control, lifeguarding, or inspection for quality control since in these jobs a drop in attention could have large detrimental consequences. In the current study, we investigated if it is possible to improve sustained attention using brain stimulation. Specifically, we tested whether transcranial electrical stimulation can prevent or reduce the time-dependent vigilance decrement.

According to Stuss et al., sustained attention relies on four distinct processes [3,4]. Activation of the task-relevant schema has to be monitored (1) and if activity drops below a certain threshold, activity of the task-relevant schema has to be reactivated (2). In addition, to reduce interference, conflicting schemata have to be inhibited (3). Finally, a higher-order process monitors task performance and takes action if performance drops (4) (through reactivation of the task-relevant schema and/or inhibition of conflicting task schemata).

The question then arises which of these processes fails during the vigilance decrement. Theoretical frameworks have attributed the time-dependent drop in performance to three (not mutually exclusive) causes: enhanced mind wandering, resource depletion, and/or reduced motivation. Some have suggested that sustained attention tasks are just not engaging enough, leading to mind-wandering [5,6]. This means that people fail to inhibit conflicting schemata and are distracted from the main task leading to the decrement in performance [7]. Others have suggested that after sustaining attention for a certain period of time, attentional resources are depleted, leading to an inability to re-activate the task-relevant schema [8]. There is much empirical evidence that supports resource depletion and this ‘mental fatigue’ account is in line with the subjective experience of the task becoming harder over time [1,9]. However, studies have also shown that social comparison or monetary rewards can improve performance [10–13]. For example, a recent study by our lab showed that the sustained attention decrement can be partly resolved by an unexpected monetary incentive [14]. This suggests that resource depletion cannot fully account for the performance decrement, but that motivation is also crucial for maintaining vigilance. This is in line with motivational control theories by Robert & Hockey [15]. Others have combined the resource depletion and motivational control accounts into hybrid models that weigh the potential reward outcome of an action against the anticipated energy expenditure [16,17]. Taken together, it seems likely that multiple factors interact to cause the vigilance decrement.

In an effort to better understand the underlying causes of the vigilance decrement, researchers have turned to neuroimaging. In a recent meta-analysis, Langner & Eickhoff (2013) identified a network of brain regions that plays a key role in maintaining sustained attention. This network comprised a large cluster spanning the presupplementary motor area (pre-SMA), midcingulate cortex, extending into more anterior medial prefrontal cortex (PFC), as well as clusters in mid- and ventrolateral PFC, anterior insula, parietal areas and subcortical structures. The authors next built a hierarchical model of how these different network nodes might interact and their putative role in sustained attention [7]. In this model, the large frontal cluster (which we will from now on refer to as medial PFC (mPFC)), plays a central role both due to its position at the top of the hierarchy and its role in performance monitoring and task-set (re-)energizing. This view corroborates a large body of evidence that links the mPFC with error monitoring and subsequent adjustments in attentional control (for review see[18]). Improper functioning of the mPFC could lead to the inability to activate the task relevant schema and thereby contribute to the vigilance decrement. Moreover, mPFC activity increases with time-on-task [7,19,20], which is thought to reflect enhanced effort to maintain attentional focus on the task [21]. In many of these studies, mPFC activity was measured through power in the theta band. Indeed, theta oscillations seem to be the lingua franca of the mPFC and are strongly linked to cognitive control processes [22]. In sum, midfrontal theta oscillations seem to play an important role in our ability to maintain sustained attention over time.

The goal of our study was to determine if electrical stimulation over the mPFC can improve the ability to sustain attention over time. To this end, we performed two experiments that used transcranial electrical stimulation. In Experiment 1, we applied transcranial *direct* current stimulation (tDCS) over the mPFC. In this technique, a small electrical current is passed between two electrodes, which is thought to depolarize (anodal stimulation) or hyperpolarize (cathodal stimulation) the underlying neural tissue [23]. We hypothesized that anodal stimulation over the mPFC would increase cortical activity in this region, resulting in improved sustained attention. This hypothesis also seems justified given previous work that has shown that anodal midfrontal tDCS can increase midfrontal theta activity [24]. Conversely, we predicted that cathodal stimulation over the mPFC would decrease cortical excitability, resulting in reduced sustained attention. In Experiment 2, we applied transcranial *alternating* current stimulation (tACS) over the mPFC. tACS is similar to tDCS, except that the direction of the current alternates at a certain frequency. This is thought to entrain neurons in the underlying neural tissue to the stimulated frequency [25]. By stimulating the mPFC in theta frequency, we expected to improve sustained attention.

## Materials and methods

### Experiment 1

#### Participants

101 healthy participants were included in the experiment and were tested in a double-blind, randomized, between-subject design. They were selected based on the following criteria; no epilepsy or (family) history of an epileptic seizure, no neurological disorders, no history of stroke or other forms of brain damage, no history of a severe concussion, no (history of) meningitis, no use of psychoactive substances, no cardiac pacemakers or other implanted medical devices, no metal anywhere in the head, no albinism, not pregnant, not recently fainted, no recent panic attack, no multiple sclerosis, no skin abnormalities on the head, and no color blindness.

All participants had normal or corrected-to-normal vision and all but three participants were right handed. Participants were instructed to abstain from alcohol, non-prescriptive medication and illicit substances 24 h prior to the experiment. If they failed to follow these instructions, they were excluded from participation.

We aimed to include at least 30 subjects per group in our final sample, i.e., double the number of subjects that previous studies included in a recent review paper (see Table 4 in [40]) and that used a between-subjects design to study the effect of transcranial electrical stimulation on sustained attention performance on average included per group (navg=15.6). Two participants decided not to continue with the experiment after the trial stimulation was applied. In addition, two participants did not complete the experiment (one was asked to leave because the participant was not doing the task, and the other participant accidently aborted the task). The remaining 97 participants (71 females) had a mean age of 22.3 years (sd = 2.7) and were randomly assigned to one of three stimulation groups: anodal (33 subjects, mean age 21.9 years, sd 2.9, 23 females), cathodal (34 subjects, mean age 22.6 years, sd 2.3, 25 females), or sham (30 subjects, mean age 22.4 years, sd 2.9, 23 females).

The protocol was approved by the Social Sciences ethical committee of the University of Amsterdam. The methods were carried out in accordance with the relevant guidelines and regulations. All participants gave written informed consent in accordance with the Declaration of Helsinki and received course credits or a monetary reward in exchange for participation.

#### Paradigm

Participants performed a sustained attention task adapted from MacLean et al. (2009). In this task (**Figure 1**), participants had to discriminate between a target (short line/33% of stimuli) and non-target (long line/67% of stimuli) and respond to the target stimuli by clicking the left mouse button with their right index finger. Throughout the experiment, participants were told to fixate on a yellow fixation dot in the middle of the screen. Between stimuli, a grey mask, composed of stacked short lines, was presented to prevent an after-image of the stimuli, which would otherwise allow participants to easily compare line lengths across trials. This mask changed on each presentation to prevent participants from comparing stimulus line length to the mask. Specifically, on each presentation, the lines that comprised the mask were vertically repositioned by a random amount (−4 to +4 pixels). Stimuli were on the screen for 150ms, while the interstimulus interval (during which the mask was presented) was randomly chosen on each trial between 1350 and 2350ms.

**Figure 1.**
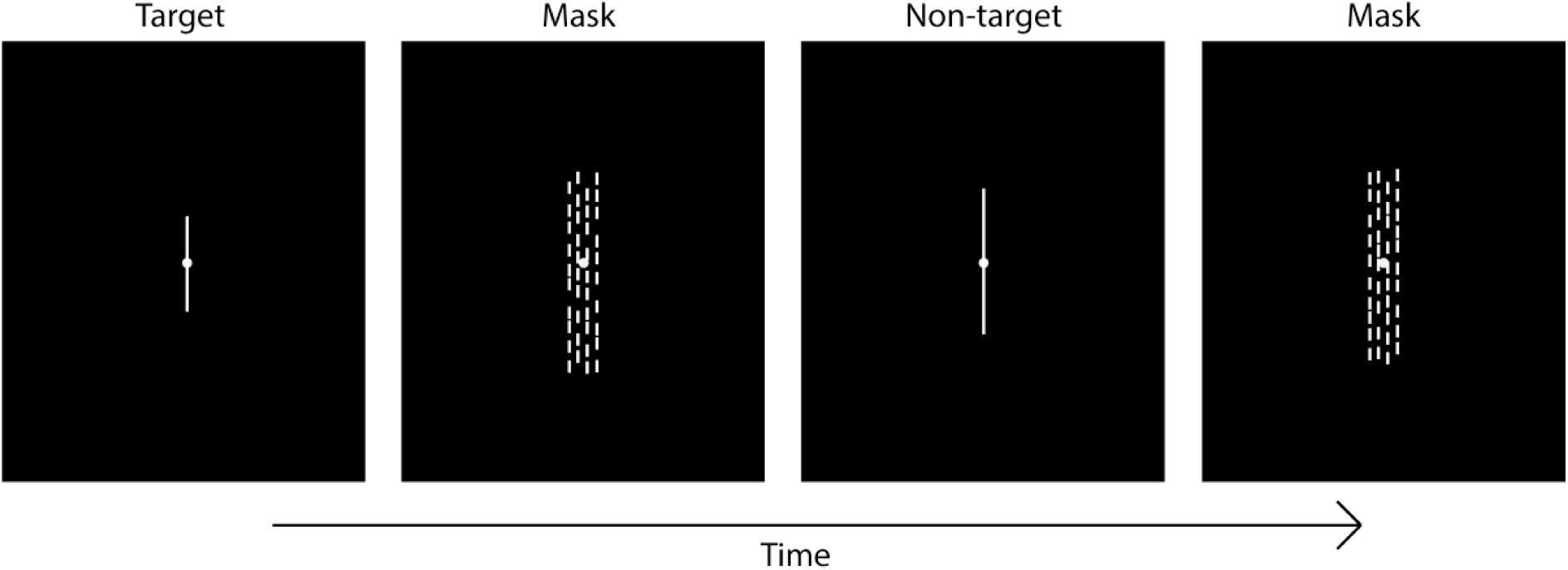
In both experiments, participants performed a sustained attention task in which they had to respond to an infrequent target stimulus (short line), and withhold responses to the non-target (long line). Presentation of stimuli (150ms) was interleaved with the presentation of masks (variable time interval, see main text).

Visual stimuli were presented using Presentation software (Neurobehavioral Systems) on a 24-inch monitor with a 1920 x 1080 resolution. Non-targets and targets had a width of 2 pixels. The height of non-targets (long line) was consistent across participants (122 pixels), while the height of the target (short line) was defined individually to balance task difficulty across participants (average=93.4 pixels, SD=13.0, range:20-112).

Individual target line length was determined with a Parameter Estimation by Sequential Testing (PEST) procedure [26,27]. During this calibration procedure, participants performed the sustained attention task described above while the length of the target line was adaptively changed according to a 3-down, 1-up staircase until performance stabilized at 80% accuracy. To help participants to learn the task, auditory feedback was provided following correct identification of a target and following a miss or a false alarm.

#### Study design

The study followed a double blind randomized between-participants design. At the start of the session, participants were checked for exclusion criteria and asked about drug and alcohol use in the last 24h. Next, participants were seated 57 cm from the monitor, received task instructions and performed the PEST procedure. Completion of the PEST procedure lasted approximately 10 minutes, but exact length varied between subjects due to the dynamic stopping rule used by the PEST titration. Upon completion of the PEST, stimulation electrodes were attached and participants were informed about any possible side effects that could occur. Participants then received one minute of trial stimulation (5 seconds ramp up, 60 seconds stimulation, 5 seconds ramp down). Participants were asked about their experience both during and after trial stimulation. If a subject reported uncomfortable side effects and/or impedance was above 25 mΩ, the set-up was adjusted to lower impedance. In this case, trial stimulation was applied again to ensure that participants felt comfortable with the stimulation. After trial stimulation, participants performed the sustained attention task for a total of 60 minutes divided over three blocks of 20 minutes. In the second of these 20-min blocks, stimulation was applied. This allowed us to examine if tDCS could revert or prevent the vigilance decrement from further developing. Before the start of each block, and at the end of the experiment, participants answered two questions about how motivated they were and how much aversion they felt toward the task by moving an arrow over an 1100 pixels wide scale by scrolling the mouse wheel. Stimulation electrodes were removed after the second 20-minute block. After the experiment ended, the participants filled out a questionnaire about possible side effects of the stimulation, as well as the perceived effect of stimulation on task performance. Side effects were scored on a linear scale from 0 to 100, with a score of zero representing ‘not at all applicable to me’ and a score of 100 representing ‘completely applicable to me’. The perceived effect of stimulation on task performance was rated as worse, similar or better performance compared to the blocks without stimulation. Ratings for 10 possible side effects were provided: headache, neck pain, nausea, muscle contractions, stinging sensations, burning sensations, tiredness, mood changes, uncomfortable feeling, and phosphenes.

#### tDCS

tDCS (NeuroConn DC-Stimulator MR, NeuroConn, Germany) was administered via two rubber electrodes. Electrodes were placed in saline soaked sponges and attached to the head via straps and tape. One electrode (9 cm2) was placed between electrode locations FCz and Cz on the 10-20 international EEG system and a second electrode (35 cm2) was placed on the left cheek [28]. Stimulation intensity was set to 1 mA (current density of 0.11 mA/cm^2^ at mPFC electrode). During the anodal stimulation condition, the anode was placed at Cz/FCz and the cathode at the cheek, while the opposite was true for the cathode condition. Stimulation was ramped up for 60 seconds, and after the 20-minute stimulation period, ramped down again for 60 seconds. In the sham group, half of the participants received ‘anodal’ sham stimulation, while the other half received ‘cathodal’ sham stimulation. In this control condition, stimulation was ramped up for 60 seconds and then ramped down again over 60 seconds. Experimenters were blinded to stimulation condition, as the stimulation settings were programmed by a colleague beforehand and they could not see the settings.

#### Data analysis

Data were analyzed using Matlab (The Mathworks, Inc.), SPSS (IBM SPSS Statistics, version 23.0) and JASP (version 0.12.2)[29]. Per 20-minute block (pre, during and post stimulation), we determined the hit rate and false alarm rate and subsequently calculated the non-parametric perceptual sensitivity measure A’ as formulated by Stanislaw and Todorov [30] (and implemented cf. [14,27,31]). A’ is based on signal detection theory and indicates a subject’s ability to discriminate between targets and non-targets. Values typically range from 0.5 (subjects cannot distinguish targets from noise) to 1 (subjects distinguish all targets from noise) [30]. In addition, we calculated the average reaction time per block.

Data were submitted to repeated measures ANOVAs with the between-subject factor stimulation condition (sham vs. cathodal vs. anodal) and the within-subject factor time (pre stimulation vs. during stimulation vs. post stimulation). Post-hoc paired t-tests were used when significant results were found. In addition, ANCOVA’s were used in order to assess whether reported side effects and target line length differed across stimulation conditions. When possible, we also conducted Bayesian variants of these statistical tests, to quantify the strength of evidence for or against an effect. We report the BF10 when the alternative hypothesis was tested, and BF01 when the null-hypothesis was tested. BF values reported for interaction effects indicate the extent to which there is more/less evidence for the model that includes the interaction effect (next to the main effects) vs. the model that only includes the main effects.

### Experiment 2

#### Participants

The methods were the same as in Experiment 1 except for the following changes. 55 healthy participants were included in the experiment and were tested in a randomized, within-subject design. In addition to the exclusion criteria mentioned for Experiment 1, we also screened for spondylosis, scoliosis, arthritis and frequent occurrences of dizziness or headaches. The sample size tested was based on previous studies using a within-subjects design to study the effects of transcranial electrical stimulation on sustained attention task performance included a recent review (see Table 4 in [40]) and chosen to be twice as large as the average sample size (n_avg_=27.6) used in these previous studies.

All participants were right-handed and had normal or corrected-to-normal vision. Three participants decided not to continue with the experiment after the trial stimulation was applied and three participants were eliminated due to technical reasons (stimulation was automatically aborted at some point during the stimulation block due to high impedance levels). One participant drank alcohol within the 24h before the experiment and was therefore excluded. Finally, one participant dropped out after the first session. The remaining 47 participants (34 female) had a mean age of 21.5 years (SD=2.6).

#### Paradigm

The paradigm was adapted in two ways. First, the target/non-target ratio was adapted, so that now 25% of stimuli was a target (compared to 33% in Experiment 1). Second, the inter-stimulus interval lasted a minimum of 1550ms and a maximum of 2150ms (compared to 1350 – 2350ms in Experiment 1). Again, the height of the target (short line) was defined individually, on each session, to balance task difficulty across participants (average over two sessions=95.1 pixels, SD =8.3, range: 55-111).

#### Study design

In contrast to the first experiment, we used a within-subject design in the second experiment to enhance statistical power. Participants participated in two sessions separated by 1 week (except for 6 participants in which the sessions were separated somewhere between 6 and 15 days). The two sessions were similar except for the frequency of the applied tACS stimulation. The order of the stimulation condition was randomized and both participants and the experimenter were blind to the stimulation condition (double blind design). The general outline of the sessions was similar to the one in the first experiment with the exception of the following: 1) trial stimulation consisted of one minute of alpha stimulation (10 seconds ramp up, 30 seconds stimulation, 10 seconds ramp down), 2) participants performed the sustained attention task for a total of 50 minutes (15 minutes before and after stimulation, and 20 minutes during stimulation), 3) side effects were scored on a five-point scale with a score of one representing ‘not at all applicable to me’ and a score of five representing ‘completely applicable to me’. In addition, we tried to minimize breaks during the task as much as possible. Therefore, the questions about motivation and aversion were eliminated from the design and we did not remove the stimulation electrodes until after the end of the experiment. Thus, there were no pauses during the experiment except for the time required to start stimulation in between block one and two, which lasted approximately one minute. No interaction with the participants took place during this minute to minimize the duration of the task break. Participants were seated approximately 65 cm from the screen.

#### tACS

1mA tACS (NeuroConn DC-Stimulator MR, NeuroConn, Germany) was administered via three rubber electrodes. Electrodes were placed inside saline soaked sponges and attached to the head via straps and tape. The target electrode (9 cm^2^) was placed between electrode locations FCz and Cz on the 10-20 international EEG system as in Experiment 1. Two reference electrodes (35 cm^2^) were placed on the cheeks. This set-up has previously been used by van Driel et al. [28]and is aimed to direct current flow through the mPFC. Stimulation intensity ranged between -1 and +1 mA and was set to sinusoidal (current density of 0.11 mA/cm^2^ at mPFC electrode). Fading in and out of the stimulation lasted for ten seconds each.

For the active condition, stimulation frequency was set to 4 Hz (theta band). During the control condition and trial stimulation, 10 Hz (alpha band) tACS was applied. Alpha band stimulation instead of sham was used for the control condition because tACS is often accompanied by phosphenes (visual disturbances) and cutaneous sensations under the electrodes, which possibly enables participants to differentiate between active stimulation and the sham condition [32]. A control condition producing similar side effects is thus needed to ensure that the active and control sessions are as similar as possible. The current study used alpha band tACS because van Driel et al. (2015), who also used theta band stimulation as the active condition and alpha band stimulation as the control condition, reported that theta and alpha tACS resulted in similar side effects. Furthermore, stimulation with higher frequency bands, such as beta frequencies, has been associated with higher phosphene intensity compared to theta band tACS [33]. Finally, while lapses in sustained attention have been associated in some studies with increased alpha activity, these changes in alpha activity occur over posterior scalp regions [e.g., [34]], i.e., not over the anterior scalp regions at which we applied tACS. We therefore did not expect midfrontal alpha tACS to directly modulate sustained attention performance.

#### Data analysis

A’ and reaction time were calculated as in Experiment 1. Next, data were submitted to repeated measures ANOVAs with the within-subject factors Stimulation Frequency (alpha vs. theta) and Time (pre, during and post stimulation). Post-hoc paired t-tests were used when significant results were found. In addition, Wilcoxon Signed Rank tests were used to assess whether side effects differed between stimulation frequencies. Because, indeed, some self-reported side effects were significantly different (see Results), we re-analyzed A’ using a linear mixed model, allowing us to include these side effects per stimulation frequency as a covariate. The model tested for main effects of stimulation frequency and time. In addition, it tested for main effects of tingling sensation and light flashes. The model also included a time x stimulation frequency interaction.

Lastly, a paired samples t-test was used to assess whether target line length differed significantly between stimulation frequencies.

## Results

### Experiment 1

We found no significant differences in target line length between the different stimulation groups (F(94)=2.269, p=0.109, BF_01_ = 1.737), indicating that task difficulty was similar across groups (anodal, cathodal and sham stimulation).

In line with previous studies, perceptual sensitivity, as measured by A’, declined with time on task (main effect of Time: F(1.8, 167.2)=16.872, p<0.001, BF_10_=109645.144) (Figure 2, left). However, there was no Time x Stimulation Condition interaction (F(4,188)=0.325, p=0.861, BF_10_ =0.029), nor a main effect of Stimulation Condition (F(2,94)=1.534, p=0.221, BF_10_=0.604). Thus, in contrast to our main prediction, perceptual sensitivity was not modulated by anodal or cathodal tDCS over the mPFC.

**Figure 2.**
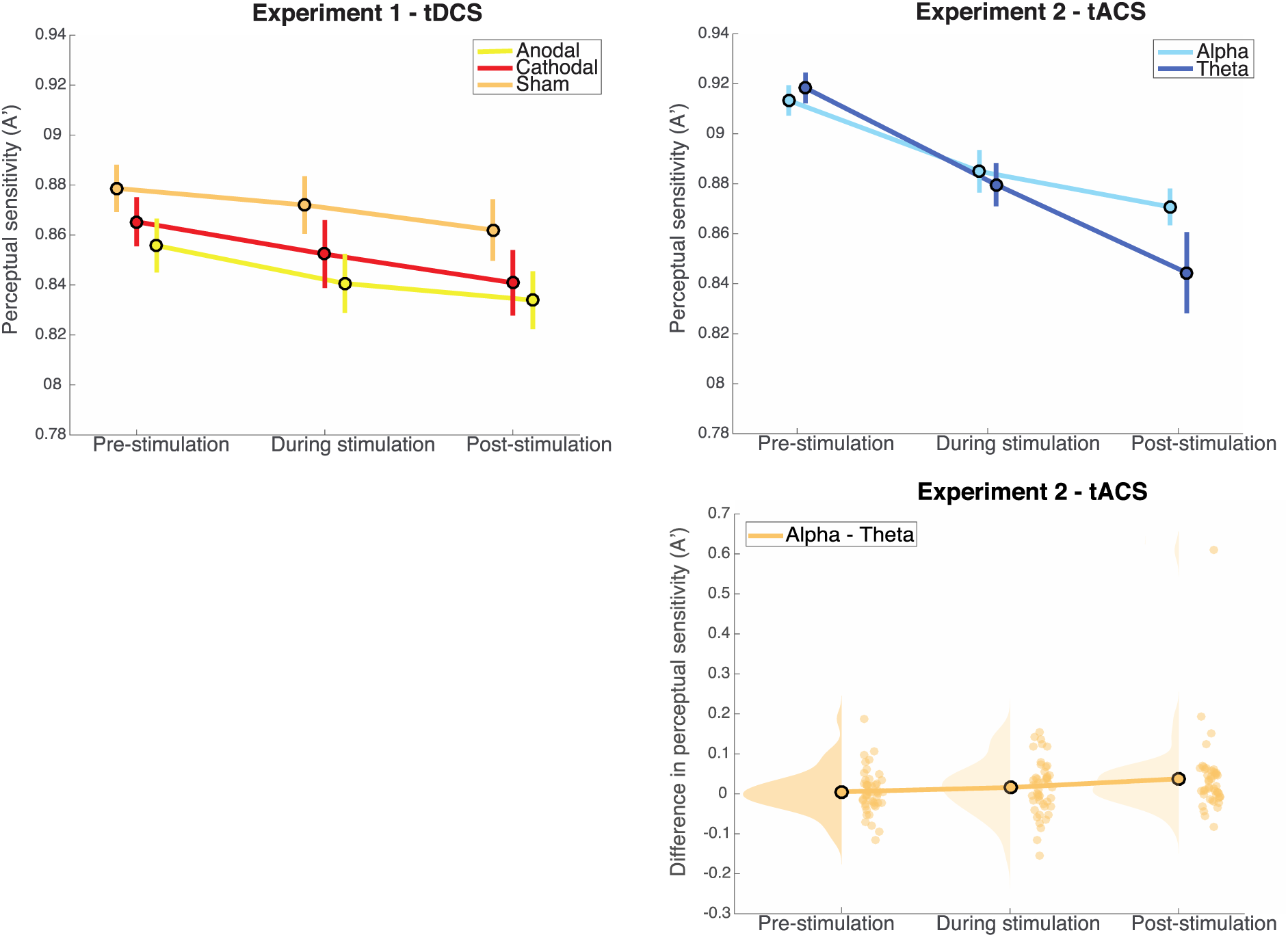
In both experiments, a robust vigilance decrement was observed, as perceptual sensitivity (A’) decreased significantly over time. In Experiment 1 (left), we found no significant effect of stimulation on the vigilance decrement. In experiment 2 (right), we found a significant Time x Stimulation frequency effect, such that perceptual sensitivity decreased more over time after theta stimulation compared to alpha stimulation. The upper right plot shows the data for alpha and theta stimulation separately. For the lower right plot we subtracted data from the theta condition from the data from the alpha condition for each individual. Next, we plotted the mean difference as well as the distribution and the individual data points [35].

For RT, no effects were found (main effect of Time: F(2,188)=1.469, p=0.233, BF10 =0.144, main effect of Stimulation Condition (F(2,94)=1.161, p=0.318, BF_10_ =0.520), Time x Stimulation Condition interaction (F(4,188)=0.681, p=0.606, BF_10_ =0.056)).

The three different stimulation conditions led to a significant difference in self-reported headache. Specifically, a series of one way ANOVAs showed a main effect of Stimulation Condition on the amount of headache (F(2,94)=4.926, p=0.009, BF_01_ = 0.215), which was driven by the fact that participants that received anodal stimulation reported a higher amount of headache compared to the other participants (anodal vs. cathodal: t(41.0)=2.394, p=0.021, BF_01_ = 0.348; anodal vs. sham: t(41.1)=2.356, p=0.023, BF_01_ = 0.457; sham vs. cathodal: t(62)=0.062, p=0.951, BF_01_ = 3.903)(median values per stimulation condition: anodal: 3; cathodal: 0; sham: 0))(mean values per stimulation condition: anodal: 12.3; cathodal: 4.1; sham: 4.2). It should be noted that when correcting for multiple comparisons (we ran an ANOVA for each of the ten self-reported side effects), the effect of stimulation condition on headache is not significant. In the anodal group, 12 participants thought their performance worsened during the stimulation block, 11 participants thought their performance remained the same during the stimulation block, and 10 participants thought their performance improved during the stimulation block. For the cathodal group and sham group, the distributions were as follows: cathodal: 13 (improved); 12 (same); 8 (worsened); 1(data missing), sham: 7 (improved); 13 (same); 10 (worsened); 1 (data missing).

### Experiment 2

We found no significant differences in target line length between the different stimulation frequencies sessions (t(46)=0.264, p=0.793, BF_01_=6.110), indicating that task difficulty was similar across conditions.

Again, perceptual sensitivity, as indexed by A’, declined with time on task (main effect of Time: F(1.4, 65.9=40.264, p<0.001, BF_10_ = 3.163e+10). Consistent with our hypothesis, we found a significant Time x Stimulation Frequency interaction (F(1.5, 71.1)=5.435, p=0.011)(Figure 2, right). However, in contrast to our prediction, this interaction was driven by the fact that performance declined more in the theta stimulation condition than in the alpha stimulation condition in particular after the stimulation had ended. Yet, the post-hoc t-test did not reach significance (t(46)=1.891, p=0.065, BF_10_=0.813), although numerically, perceptual sensitivity was lower after theta stimulation than after alpha stimulation. Note also that while the frequentist analysis indicated a significant interaction between Time and Stimulation Frequency, a Bayesian repeated measures ANOVA indicated that the data are 0.565 times less likely under the model that includes the interaction effect (BF_10_ =6.9e+9) than under the model that only includes the main effects of Time and Stimulation Frequency (BF_10_ =1.221e+10). Thus, the statistical evidence for a differential effect of stimulation frequency on sustained attention task performance over time was weak. (Note that the outlier in the bottom right plot is not driving the results, but is actually reducing the significance of the effect (F-value: 5.4 vs. 5.5; p-value: 0.011 vs. 0.006; BF_10_ = 0.565 vs. BF_10_=0.556).)

For RT, we found a main effect of Time (F(1.7, 78.7)=19.280, p<0.001, BF_10_=4993.764), but no main effect of Stimulation Frequency (F(1,46)=0.134, p=0.716, BF_10_=0.149), nor an interaction (F(1.6,74.7)=1.674, p=0.193, BF_10_ =0.167).

In terms of side effects, we also found a difference between the stimulation conditions. Specifically, participants perceived more tingling sensations (Z=-4.053, p<0.001, BF_01_ = 5.341e-4) and more phosphenes (Z=-4.981, p<0.001, BF_01_ = 2.623e-7) during alpha stimulation. This could confound our findings on perceptual sensitivity. In an attempt to control for these potentially confounding side effects, we performed another analysis, in which we tested for a Time x Stimulation Frequency interaction on perceptual sensitivity using these two side effects as a covariate. Again, we found a main effect of Time (F(2,230)=33.587, p<0.001), indicative of a decrease in performance over time, but now the Time x Stimulation Frequency interaction did not reach significance (F(2,230)=2.508, p=0.084).

## Discussion

Sustained attention is crucial in many everyday life and professional settings that require continuous monitoring to detect rare and difficult to predict events, such as when driving a car or in air traffic control. In two experiments with relatively large samples sizes (n=97 and n=47, respectively), we aimed to improve people’s ability to sustain attention over time using transcranial electrical stimulation over mPFC, a key region involved in sustained attention [7]. However, in neither experiment, we found convincing evidence that electrical stimulation over mid-frontal cortex can enhance sustained attention. These findings contribute to a growing body of studies that report no effect of electrical brain stimulation on behavior [36,37], but contrast with studies that did observe positive effects of electrical stimulation on sustained attention [38,39]. Below, we discuss our findings in greater detail with respect to the current literature, and discuss potential explanations and necessary follow-up studies.

In Experiment 1, neither anodal nor cathodal tDCS over mPFC modulated sustained attention compared to sham tDCS. It is possible that our stimulation did not reach the mPFC. Yet, it has previously been shown that 1mA anodal tDCS over midfrontal cortex can increase frontal theta activity, albeit transiently [24]. Nonetheless, in that study, no effects were observed on a subsequent sustained attention task. Our findings extend this work by showing, using a much larger sample, that midfrontal anodal tDCS concurrent with a sustained attention task is also not associated with performance improvements. In previous studies looking at the effect of tDCS on sustained attention, it was found that tDCS over bilateral dorsolateral prefrontal cortex (DLPFC) could prevent the vigilance decrement [38,39]. However, replication of these effects are warranted since some of the effects ascribed to tDCS were already apparent before tDCS stimulation started [38]. Also, correction to the baseline block, as was applied in these studies, can lead to artificially induced effects, by adding baseline differences to experimental effects. Alternatively, it is possible that we did not choose the most optimal settings in the design of our experiment. We used a different montage than a previous study that reported tDCS effects on sustained attention with bilateral frontal stimulation (although in the presence of baseline differences) [39]. Although 1mA is a common current strength used for tDCS in attention studies (see for example a recent review by Reteig et al. [40]), the effects of tDCS may vary as a result of the applied stimulation intensity [41], though not necessarily in a linear fashion [42]. Another reason for the null findings could be the fact that we used a between-subject design. tDCS effects are generally subtle and might therefore surface only in a within-subject design which is more sensitive because it takes out individual variance in baseline task performance and the degree of vigilance decrement over time (although we tried to control for the first with the staircase procedure). However, compared to many tDCS studies that previously reported effects on attentional performance using a between-subjects design [40], our sample size was relatively large with 30 to 34 participants in each stimulation group. It should also be noted that the effects of tDCS in general are debated [43]. As we pointed out in an earlier review, the wide range in stimulation parameters that is used in tDCS studies of attention and individual differences in the effects of tDCS might contribute to the failure to find (consistent) effects [40]. Also, the neural effects of tDCS itself may be smaller than assumed, as it was recently shown that the strength of the electric field in the brain is at the lower bound for it to be physiologically effective [44,45]. Notably, a recent study that applied cathodal tDCS over left lateral prefrontal cortex found that this increased mind wandering during a sustained attention to response task, but only at 2mA [53]. Future work is necessary to determine if higher stimulation strength could similarly affect performance on sustained attention tasks, that only rarely require a response, as used in our study. To summarize, using 1mA anodal tDCS over mPFC we were not able to improve sustained attention.

In Experiment 2, using a within-subject design with 47 participants, we found effects of tACS on the vigilance decrement. However, we have to be cautious about the interpretation of these results. First, in contrast to our expectations, participants’ vigilance decrement became worse after theta stimulation compared to alpha stimulation. As explained in the introduction, we anticipated that theta stimulation would improve performance based on previous studies showing that theta oscillations originating from the mPFC are involved in performance monitoring and task-set (re-)energizing [22]. It is possible that we interfered with an already optimally functioning system (in healthy individuals) and therefore found the opposite of what we anticipated. Also, we did not use individually determined theta frequency, but stimulated everyone at 4 Hz. It has been suggested that effects of tACS might be strongest when stimulation happens in sync with the individual intrinsic frequency [46,47]. Perhaps by entraining mPFC neurons to a frequency that did not match their intrinsic frequency in some of our participants, we disrupted the system, rather than helped it. Yet, some studies have also reported no effects of midfrontal theta tACS, albeit on a working memory task[48]. Alternatively, it is hence also possible that theta stimulation had no effect, but that alpha stimulation improved performance. Indeed, in a recent study, Clayton et al. found that alpha stimulation over occipital cortex stabilized performance on a visual attention task [49]. The researchers speculated that the alpha stimulation may have prevented top-down reorienting signals from changing the attentional state of the visual system. Alpha stimulation may similarly have stabilized the monitoring state of mPFC. Indeed, sustained attention has not only been associated with oscillations in the theta frequency, but also in the alpha and gamma range [21,50,51]. Frontal alpha-band oscillations may also play a role in cognitive control (e.g., [52]). Yet, without the presence of a sham condition, we cannot dissociate whether the effects that we found are driven by effects in the alpha stimulation condition, the theta stimulation condition, or a combination of both. However, as described in the methods section, including a sham condition has its own disadvantages. Moreover, it should be noted that the results from a Bayesian repeated measures ANOVA did not support the frequentist result of an effect of stimulation frequency on sustained attention task performance over time, as this analysis suggested that the data were almost two times more likely under the model that did not include the interaction effect.

Another difficulty in the interpretation of the obtained results is the fact that we found differences in side effects between the two stimulation frequencies. During alpha stimulation, subjects experienced more tingling sensations and more phosphenes compared to during theta stimulation. It could be argued that these side effects mainly occurred during the block of stimulation and do not extend to the block after stimulation, in which the difference between the stimulation conditions seems to arise. Also, it seems counterintuitive that an increase in side effects, would lead to better rather than worse perceptual sensitivity. One possibility is that theta-induced phosphenes, albeit less frequently experienced, were experienced as much more unpleasant or disruptive than alpha-induced phosphenes. Moreover, we cannot exclude the possibility that experiencing side effects made subjects more alert or strongly believe that they were in an active stimulation condition, thereby increasing their motivation to do well. When trying to control for these differences, the Time x Stimulation Frequency interaction no longer reached significance. This suggests that at least part of the frequency-dependent tACS effects on the vigilance decrement can be explained by the differences in side effects between the stimulation frequencies. More work is necessary to better establish at what stimulation intensity side effects, such as phosphenes, may arise at different stimulation frequencies, as they are clearly confounding factors. In sum, although we do find frequency-dependent tACS effects, with the current setup we cannot determine whether participants improved or worsened by the stimulation, and they could simply reflect non-specific side effects. Further research including other control conditions, for example stimulation over another brain region, that evoke similar side effects, is needed to determine the direction of the observed effects and the role of midfrontal theta oscillations in sustained attention.

## Conclusions

To conclude, across the two studies we were not able to find a convincing method to prevent or reduce the vigilant decrement. More research that combines brain stimulation with neuroimaging, is needed to determine if and how electrical stimulation over mPFC may affect brain functioning and sustained attention.

## Competing interests

The authors declare no competing interests.

## Acknowledgements

This research was funded by a Marie Curie European Fellowship grant to MS and a VIDI from the Netherlands Organization for Scientific Research (NWO) to HAS.

## Author contributions

MRS, IGS, HAS conceived and designed the experiments; MRS, IGS, MRG, FG, DK, FSS, AV collected and analyzed the data; MRS, IGS, and HAS interpreted the data and wrote the manuscript. All authors reviewed the manuscript.

## Data availability

Data and analysis scripts will be made available after acceptance for publication via the University of Amsterdam Figshare system (https://uvaauas.figshare.com/).

## Notes

### Competing Interest Statement

The authors have declared no competing interest.

